# Deep learning models of perceptual expertise support a domain-specific account

**DOI:** 10.1101/2022.12.01.518342

**Authors:** Galit Yovel, Idan Grosbard, Naphtali Abudarham

## Abstract

Perceptual expertise is an acquired skill that enables fine discrimination of members of a homogenous category. The question of whether perceptual expertise is mediated by general-expert or domain-specific processing mechanisms has been hotly debated for decades in human behavioral and neuroimaging studies. To decide between these two hypotheses, most studies examined whether expertise for different domains is mediated by the same mechanisms used for faces, for which most humans are expert. Here we used deep convolutional neural networks (DCNNs) to test whether perceptual expertise is best achieved by computations that are optimized for face or object classification. We re-trained a face-trained and an object-trained DCNNs to classify birds at the sub-ordinate or individual-level of categorization. The face-trained DCNN required deeper retraining to achieve the same level of performance for bird classification as an object-trained DCNN. These findings indicate that classification at the subordinate- or individual-level of categorization does not transfer well between domains. Thus, fine-grained classification is best achieved by using domain-specific rather than domain-general computations.

## Introduction

Perceptual expertise refers to the exceptional ability to identify members of a homogenous category. This acquired skill is achieved by extensive experience in classifying stimuli of a specific category, such as birds or cars at the subordinate or individual-level of categorization ^1–4^. Faces are the only category for which most humans are expert. The ability to classify faces to different individuals is acquired thorough the extensive social and perceptual experience that humans have with people. Thus, a major question that has been hotly debated for over three and a half decades ^5^, is whether perceptual expertise is mediated by domain-specific or general-expert processing mechanisms (for reviews see, Bukach et al., 2006; McKone et al., 2007). According to the domain-specific hypothesis, expertise for faces is mediated by mechanisms that are specific to the domain of faces and is not used for expertise for other domains, such as birds or cars ^6,8–10^. Conversely, according to the general-expertise hypothesis, classification of objects of expertise, including faces, is mediated by a general mechanism for subordinate level classification^7,11,12^.

To decide between these two hypotheses, previous studies examined whether real-life or lab-based experts show the same behavioral and neural face-selective markers for their objects of expertise. These face-selective markers typically included the face inversion effect (i.e., disproportionally lower performance for inverted than upright faces) ^5,8^, holistic/configural processing ^8,13,14^ and/or the face-selective neural response in the fusiform gyrus ^9,15,16^. For example, studies examined whether bird or car experts show a stronger response in the fusiform face area ^9,15,16^, or a disproportionally large inversion effect, to their domain of expertise. Whereas some studies reported that objects of expertise activate face-selective brain areas ^11,16^, other studies showed that responses to objects of expertise are not limited to face-selective brain areas but are found in multiple cortical regions ^15,17^. Similarly, studies that examined the inversion effect also yielded inconsistent findings (for review see, McKone et al., 2007), with some reporting a large inversion effect for objects of expertise that was similar to faces ^5,18^, and others reporting a smaller inversion effect for objects of expertise than for faces ^8,13,19–21^. Therefore, currently there is no agreement among researchers on whether the data support the domain-specific or general-expertise hypothesis.

In the current study, we used deep learning algorithms to test the computational plausibility of the two hypotheses. Deep convolutional neural networks (DCNNs) are brain-inspired algorithms that reach human-level performance and generate human-like representations for objects and faces ^22–25^. These algorithms can be used to model perceptual expertise by training them to classify images from different domains at different levels of categorization. To decide between the general-expertise hypothesis and domain-specific hypothesis, in the current study, we re-trained (i.e., fine-tuned) DCNNs to classify bird stimuli as the new domain of expertise. Similar to faces, birds are an animate category, which has been used before in human studies to decide between the two hypotheses (e.g., Gauthier et al., 2000). We selected two data sets of bird stimuli: one dataset that includes different bird species – this parallels bird experts who can discriminate bird species at the subordinate level of classification, and a second dataset of individual birds from a specific species of Sociable Weavers ^26^ that, similar to faces, are classified at the individual level of categorization.

To examine whether different domains of expertise are processed by the same or different computations, we re-trained (fine-tuned) an expert DCNN, which is optimized for classification of faces at the individual level of categorization, and a non-expert DCNN, which is optimized for classification of objects at the basic-level of categorization, to classify a new domain at the subordinate or individual-level of categorization (Figure 1).

**Figure 1:**
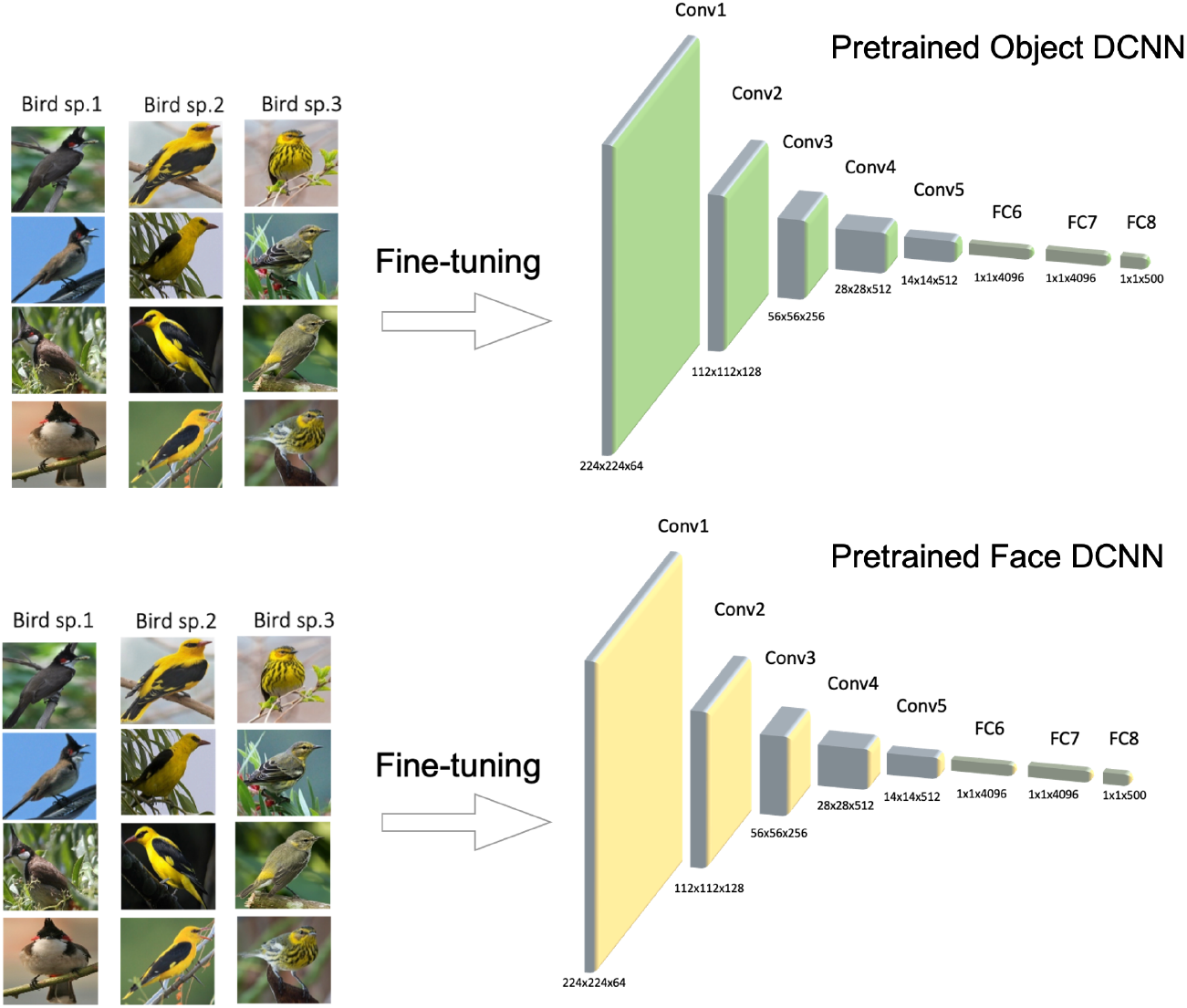
Two DCNNs were pre-trained, one for classification of inanimate objects (nonexpert), and one for classification of faces (expert), using an identical architecture (VGG16) a similar training protocol and the same number of images. We then re-trained (fine-tuned) each of the pre-trained DCNNs to perform a bird classification task. We generated 7 different fine-tuned DCNNs for the 8-layer network, by re-training an increasing number of layers, starting with the top layer until re-training the whole network for birds. Performance for bird classification was measured based on embeddings of the bird images in the penultimate layer (FC7).

The general-expertise hypothesis predicts that a network that is expert in one domain (faces) can re-learn (transfer to) a new domain (birds) more easily than a non-expert network (objects). The domain-specific hypothesis predicts that a network that is expert in one domain (faces) will not transfer well to a new domain (birds) and therefore makes the opposite prediction (Figure 2). This can be examined by measuring the number of re-trained layers required for the face and object networks to reach the same performance level for bird classification.

**Figure 2:**
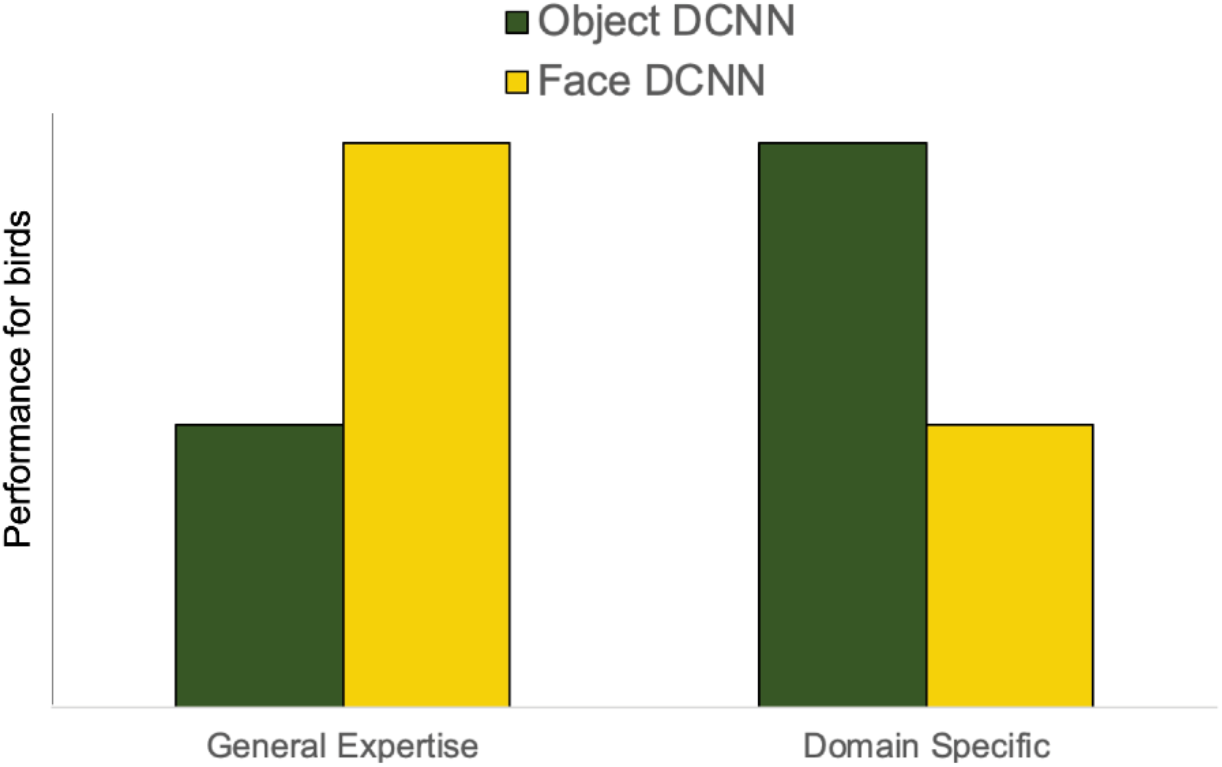
Predications of the general-expertise and domain-specific hypotheses: Performance for bird classification in face and object pre-trained DCNNs that were re-trained (fine-tuned) to classify birds. The general-expertise hypothesis predicts better performance for birds in a pre-trained face than a pre-trained object DCNN that are re-trained to classify birds. The domain-specific hypothesis makes the opposite prediction.

## Results

### Expertise for bird species

Figure 3 shows the performance for bird species classification for object and face DCNNs before retraining and following re-training of each additional layer of the networks. Performance of the object DCNN to classify birds was better than the face DCNN before retraining. The face DCNN reached performance level of the pre-trained object DCNN, only after the two fully connected layers were re-trained to classify birds. A mixed ANOVA with Pre-trained domain (Object, Face) as a between factor and Layer as a within factor, revealed a main effect of Domain (F(1,58) = 235.6, p < .001, η^2^_p_ = .80) that reflects better performance for birds after fine-tuning a pre-trained object than face DCNN. An interaction between Layer and Domain (F(7,406) = 132.25, p < .001, η^2^_p_ = .69) indicates that this advantage for the pre-trained object network is found only when the fully connected layers are fine-tuned for bird classification. As shown in Figure 3, performance for birds of the pre-trained face and object networks start to converge after fine-tuning included the convolutional layers (Conv1 to Conv5). This is in line with our recent findings that showed that these convolutional layers generate similar representations in face and object-trained networks, which diverge to different representations only in the fully connected layers ^22^.

**Figure 3:**
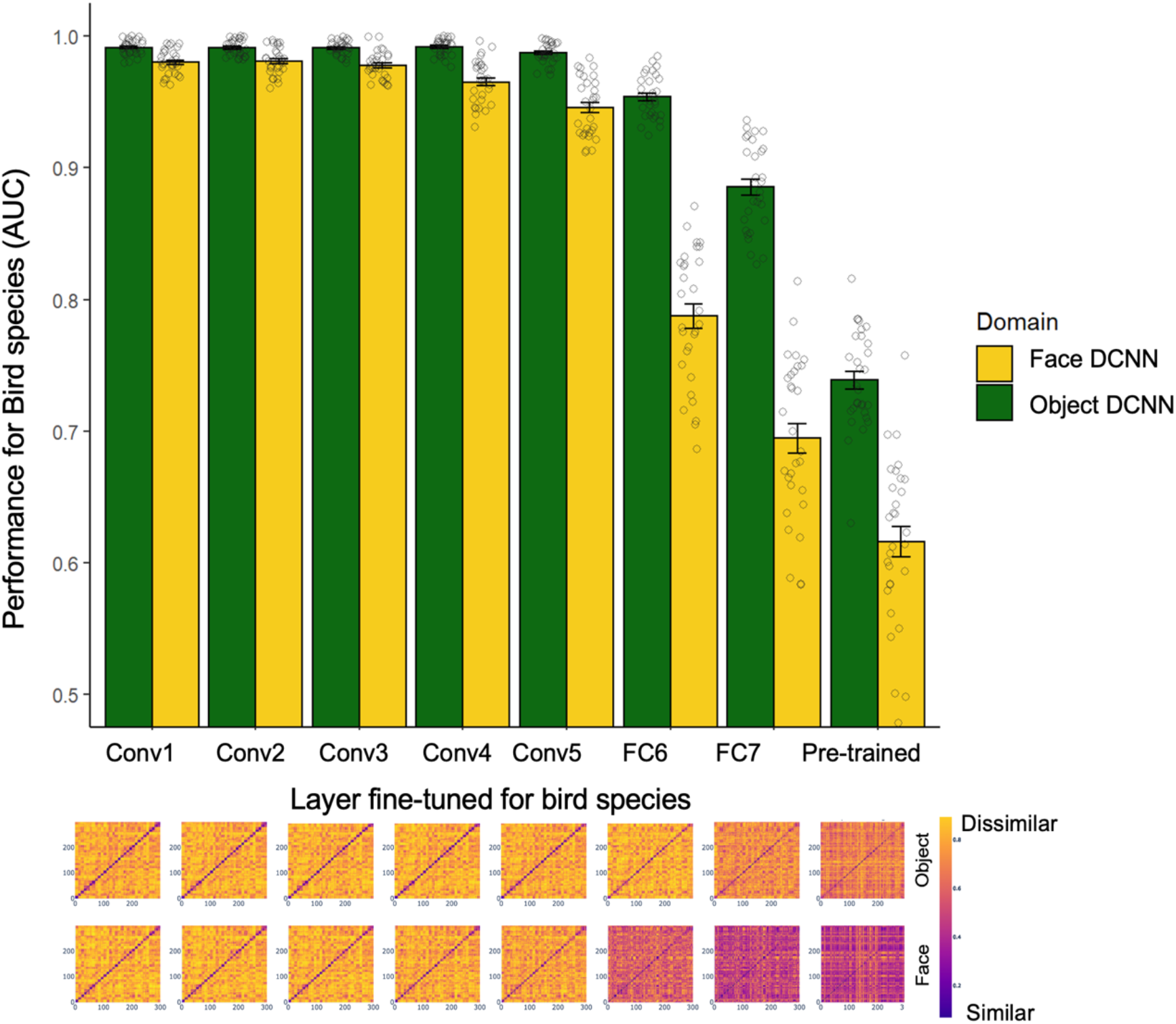
Performance for bird species for pre-trained face (yellow) and object (green) DCNNs before and after fine-tuning to classify birds. The x-axis indicates the first layer that was fine-tuned for bird species (i.e. the layers before this layer were freezed). Performance improves as the number of fine-tuned layers increases. Representation Dissimilarity Matrices (RDMs) for bird species based on their representations in the penultimate (FC7) layer in pre-trained object (upper row) and face (bottom row) networks and after fine-tuning additional layers. Better classification is indicated by a darker diagonal, which reflects a smaller distance between images of the same class and a yellower off diagonal which reflects a larger distance between images from different classes.

These results indicate that the computations used to classify objects better fit bird classification, than the computations that are used to classify faces. Notably, the object pre-trained DCNN was only trained on inanimate stimuli from the ImageNet database. This is different from standard object-trained DCNNs that typically include classes of different animals, including birds. Thus, despite the fact that the pre-trained object network was not exposed to any animate visual features, it shows better learning for a new domain of animals, indicating that training a network to classify images at the basic-level of categorization involves computations that are better generalized to classification of new domains at the sub-ordinate level.

Bird species are classified at the subordinate level of categorization, whereas faces are classified at the individual level of categorization. We therefore obtained another database of bird images that can be classified at the individual level of categorization (see Methods), and re-trained a face and an object pre-trained networks to classify individual birds (Figure 4)

**Figure 4:**
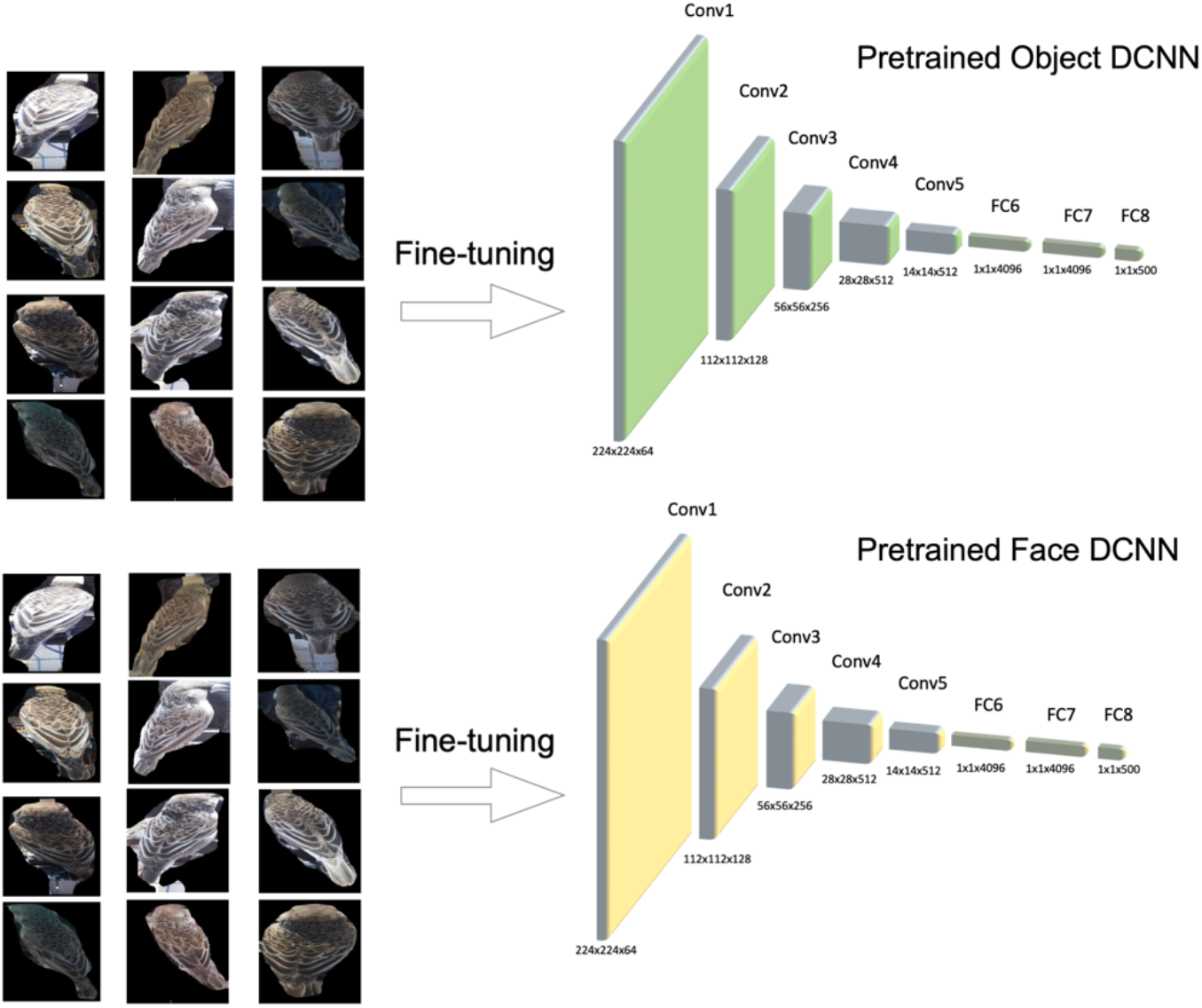
Two DCNNs were pre-trained, one for classification of inanimate objects (nonexpert), and one for classification of faces (expert), using an identical architecture (VGG16) a similar training protocol and the same number of images. We then re-trained (fine-tuned) each of the pre-trained DCNNs to perform <u>an individual bird classification task</u>. We generated 7 different fine-tuned DCNNs for the 8-layer network, by re-training an increasing number of layers, starting with the top layer until re-training the whole network for birds. Performance for bird classification was measured based on embeddings of the bird images in the penultimate layer (FC7).

### Expertise for individual birds

Figure 5 shows the performance for individual bird classification for pre-trained object and face DCNNs before re-training, and following re-training of each additional layer of the networks. Since classification of individual birds is a much more difficult task than classification of bird species, both pre-trained DCNNs are close to chance level for this task (Figure 5). Performance of the object DCNN improves after fine-tuning one fully connected layer, whereas the face DCNN requires fine tuning of two layers to reach the same performance level. A mixed ANOVA with pre-trained domain (Object, Face) as a between factor and Layer as a within factor, revealed a main effect of Domain (F(1,58) = 13.06, p < .001, η^2^_p_ = .18) that reflects better performance for individual birds after finetuning a pre-trained object than face network. An interaction between Layer and Domain (F(7,406) = 16.26, p < .001, η^2^_p_ = .22) indicates that performance for individual birds was better in the pre-trained object than face DCNNs when fine tuning the fully connected layers and merged after fine-tuning the convolutional layers for bird classification (Figure 5).

**Figure 5:**
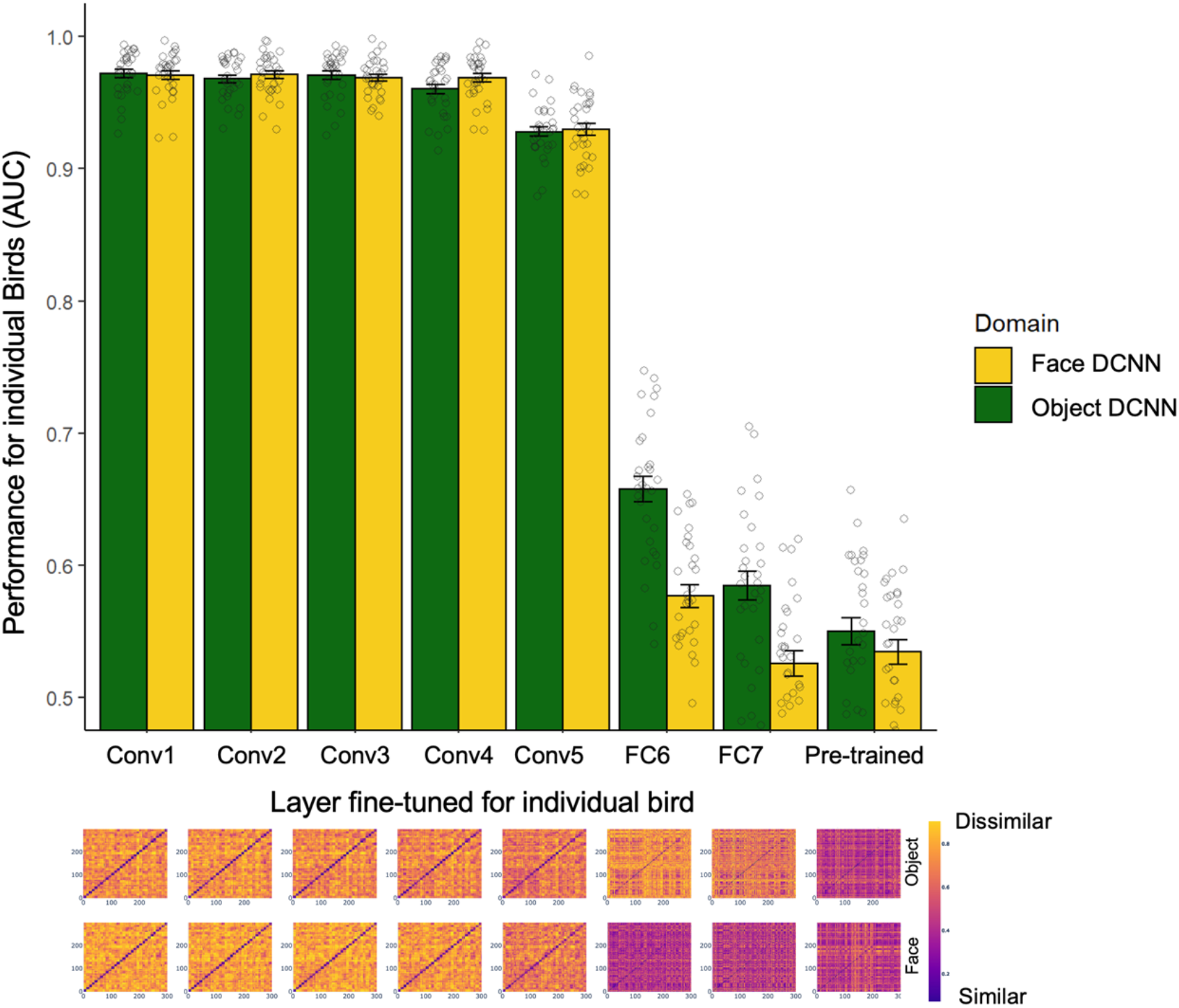
Performance for individual birds after fine-tuning each additional layer of the pre-trained face and object DCNNs, shows better performance for the pre-trained object then pre-trained face DCNN, until the two networks converge in the convolutional layers. The x-axis indicates the first layer that was fine-tuned for individual birds (i.e. the layers before this layer were freezed). The same data presented in RDMs that indicate the similarity between same and different pairs of individual birds after fine-tuning additional layers for bird classification. A darker diagonal indicates higher similarity for same than different pairs.

## Discussion

To test whether computations required for expert object classification are similar across domains, as predicted by the general-expertise hypothesis, we re-trained an object-trained (non-expert) and a face-trained (expert) deep learning networks to classify birds at the sub-ordinate or individual-level of categorization. Our results show that an expert system that is optimized to classify stimuli at the individual level of one category (faces) does not transfer well to other subordinate or individual-level categories (bird species or individual birds). In fact, it was the network that was pre-trained at basic-level classification that showed better transfer to a new domain. These findings are inconsistent with the general-expertise hypothesis, which predicts that individual-based classification uses the same computations for different domains. Instead, these findings suggest that perceptual expertise is enabled by specific training of the network to domain-specific features that are diagnostic to the learned category, and this learning does not transfer well to other categories that rely on different computations and/or features for fine-detailed classification.

The question of whether perceptual expertise is mediated by general or specialized processing mechanisms has generated numerous studies that employed multiple methods to decide between the two hypotheses (for review see, Gauthier & Bukach, 2007; McKone et al., 2007). Here we used deep learning models of object recognition, which reach human-level performance in face and object recognition, to directly test whether objects of expertise apply similar or specialized computations. But how valid are these deep learning models for human face and object recognition? Many recent studies have started addressing this question by measuring the correlations between the representations that are generated for the same set of stimuli in primate behavioral and neural measures, and the embeddings of DCNNs optimized for object and face recognition. These studies show remarkable correspondence between the representations of DCNNs optimized for object and face recognition and biological and behavioral measures of the same stimuli, accounting for a much larger proportion of variance in behavioral and neural measures than previous computational models ^24,28,29^. This correspondence goes beyond measures of performance level and show that DCNNs and humans generate similar representations and are sensitive to the same features. For example, in a recent study, Abudarham et al (2021) showed that a face-trained DCNNs is sensitive to the same view-invariant facial features that human use for face recognition. This sensitivity was significantly smaller in an object-trained DCNN, indicating that specific training with faces is needed to generate this sensitivity to view invariant facial features ^22^. They also found that the representations that are generated by a face and an object-trained networks are similar at the initial layers of both networks, but diverge at higher layers, similar to the primate visual system that diverges to a face and an object system at high level but not low-level visual cortex. These findings are also consistent with the current findings, which show that fine-tuning that includes the convolutional layers generates similar performance level for objects of expertise in the pre-trained object and face networks (Figures 4, 5).

Another human-like dissociation between face and object-trained DCNNs was recently reported by Jacob and colleagues, who showed a Thatcher illusion effect, in high-level layers of a face-trained but not an object-trained DCNN ^25^. In a Thatcher illusion, two faces in which the eyes and mouth are inverted in one face image but not the other, look more different when presented upright than inverted. Similarly, the distance between the embeddings of these face stimuli is larger when presented upright than inverted in the higher layers of a face-trained but not object-trained DCNN. A face inversion effect in a DCNN was also recently reported by Tian and colleagues ^30^. Another evidence for the specificity of the face processing system of humans and DCNNs, to the stimuli it was trained with, is the other race effect. Numerous studies in humans show better performance for own than other race faces ^31–34^, and a similar effect is also found for DCNNs ^35,36^. In both humans and DCNNs this effect depends on exposure to faces from a given race and can be alleviated if the system is trained to classify faces from the other race ^37^ Recently, Dobs and colleagues report that these human-like face effects are found in face networks that are optimized for face individuation but not in networks that are provided with the same amount of experience but classify all faces to one category ^38^.

Despite these remarkable similarities between DCNNs’ and humans’ representations, there are also differences^25,39^ that may limit the relevance of the current findings to humans. Past studies that have used shallow neural networks have found that expert networks show better transfer to other domains, in contrast to the findings reported here ^40,41^. It is possible that the shallow networks are less specifically tuned to the learned images. Indeed, shallow networks do not reach the same classification level as deep networks, which are able to reach high levels of classification for individuation of highly similar images. This may be enabled by specialized tuning to very fine details and therefore does not transfer well between domains. Overall, computational modelling can only indicate computational plausibility based on the type of computations it applies, which may or may not fit the human mind. Having said that, DCNNs are currently the only models that reach human level performance in object recognition and exhibit unprecedented similarities to the human mind.

Perceptual expertise has been often attributed to holistic/configural processing ^2,5^. Direct measures that were used to assess holistic processing are the composite face effect (Young, Hellawell, 1987), in which performance for aligned and misaligned half faces is compared and the whole-part effect ^43^ in which identification for face part within and without face context is compared. The current study did not test such or other measures of holistic processing in DCNNs, which may be further investigated in future studies. Such investigation will enable to ask whether holistic processing is indeed inherent to any type of perceptual expertise ^7,12,44^ or may be found for some domains but not others. Indeed, recent studies have not revealed strong correspondence between effects of holistic processing and performance on a face recognition task ^45^, and therefore the link between holistic processing and the ability to successfully classify stimuli in fine details is still unclear.

In summary, the current study used computational models of expert (face) and non-expert (object) recognition to decide between the general and domain-specific accounts of perceptual expertise. Our findings strongly support a domain-specific account for expert object recognition. Whereas computational models cannot provide direct evidence for the type of operation of a biological/cognitive system, the many similarities between humans and DCNNs that were already discovered, their brain-like architecture, and the full control that we have over their training diet, make them useful models to test existing hypotheses as well as design future studies to test new hypotheses and can therefore significantly advance our understanding of the human mind and brain.

## Methods

### Pre-training object and face classification DCNNs

We trained two randomly initialized VGG-16 ^46^ DCNNs, one to classify classes of objects (basic-level categorization), and the second to classify face identities (individual-level categorization) (see Figures 1 & 3).

#### Training data for pre-trained object DCNN

We randomly selected 500 classes of <u>inanimate</u> objects from the ImageNet dataset ^47^, each class consisted of 300 training images, and 50 validation images. We chose only classes of inanimate objects to avoid any similarity between the training data of the pretrained object DCNN and the bird images that were used as the new domain for re-training.

#### Training data for pre-trained face DCNN

To match the training data that was used for the object classification DCNN, we randomly selected 500 identities from the VGGFace2 dataset ^48^, and for each identity we used 300 images for training, and 50 images for validation. All face images were aligned prior to training using MTCNN^1 49^.

#### Training protocol

Both DCNNs were trained using cross-entropy loss. The networks were optimized using Stochastic Gradient Descent (SGD) with a learning rate of 0.01 and with PyTorch default parameters ^50^, for 120 epochs of 1000 batches, with batch size of 128. After 80 epochs the learning rate was reduced to 1e-3. Training images were normalized (object images were normalized using ImageNet mean=[0.485,0.456,0.406] and std=[0.229,0.224,0.225], and face images were normalized with default mean=[0.5,0.5,0.5] and std=[0.5,0.5,0.5]). Face images were first resized to 256×256 pixels. We then applied a random crop of 224×224 pixels, and employed a random horizontal flip ^51^. Object images were first resized to 224×224 pixels, followed by a random crop of scale in the range of 0.08 to 1.0 of the original image, and ratio in the range of 0.75 and 1.33. Images were then resized again to 224×224 pixels and we employed a random horizontal flip (the default PyTorch augmentation protocol).

### Fine-tuning for sub-ordinate (bird-species) and individual-level (individual-birds) classification

To assess whether perceptual expertise is generic or domain specific, we tested how well an expert and non-expert DCNN learn to classify a new domain at the sub-ordinate or individual level of categorization. To this end we re-trained (fine-tuned) the face and object pre-trained networks to classify a new domain (birds) at the sub-ordinate level of categorization (bird species), or at the individual-level of categorization (individual birds) and compared performance of the fine-tuned face and object DCNNs on the new domain. We created 7 different fine-tuned networks for each pre-trained DCNN, by updating the weights of a different number of layers, starting with the top layer and freezing all other layers^2^ and then fine-tuning additional layers one by one up to the first layer (Conv1).

#### Data for bird species fine-tuning

The bird dataset included 260 different bird species taken from <u>Kaggle</u>. Each class consisted of an average of 151 images (range 105-310). Since the number of images in each class was relatively small, for most classes (230 of 260 classes) we used 80% of the images per class for training and 20% for validation. For 30 classes with the largest number of images per class we used 50% images for training and 50% for validation.

#### Data for individual bird fine-tuning

The individual bird dataset included only 30 different individual Sociable Weavers, which were obtained from a dataset created by Ferreira et al. 2020. The dataset included an average of 801 images per bird (range: 369-900). 80% of the images in each class were used for training, and 20% for validation. The bird images were extracted from movies that were taken in captivity, from a rear view, therefore classification is not based on the faces of the birds but primarily on their feathers as shown from the back (see Fig. 2).

#### Fine-tuning protocol

Fine-tuning was done using cross-entropy loss. The networks were optimized using Stochastic Gradient Descent with a learning rate of 0.01 and with PyTorch default parameters ^50^ for 60 full epochs, with batch size of 128. After 50 epochs the learning rate was reduced to 1e-3. Training images were normalized using the pixel means and standard-deviations of each training dataset, and we used the same image augmentations that were used for the pre-trained object DCNN, with an addition of random rotations of up to +/- 40 degrees (which was found to improve performance in ^26^). For each of the 7 fine-tuned networks, a different number of layers were updated during training in a serial manner, by freezing the weights of the layers that precede the updated layers^2^ (Fig. 3).

### Measuring performance for bird species and individual-bird classification

To measure how well the object and face DCNNs classify images from the new domains of bird-species or individual-birds, before and after fine-tuning across the different layers, we performed a same/different classification task (i.e., verification). To test performance on the same/different classification task on the individual-bird images, we randomly selected 50 distinct image pairs from the validation images of each of the 30 classes of the individual birds, making 1500 (30×50) same-class pairs. In addition, we randomly selected 1500 distinct different-class pairs from the validation images. For the bird species we used the same number of same and different image pairs from the 30 classes with the largest number of images, out of the 260 classes of bird species. We generated representations of the tested images in the pre-trained and fine-tuned DCNNs, by running the images through the DCNNs and obtaining the embedding from the penultimate layer (layer FC7, see Fig. 3).

To test the performance of each DCNN on the same/different classification task, we randomly divided the 1500 same-class image pairs, and the 1500 different-class image pairs, to 30 batches. Then, for each batch, we measured the cosine similarity between the embeddings of the images in the penultimate layer for each image-pair, and calculated Receiver Operator Curves (ROCs) as well Areas Under Curves (AUCs) for true and false same-class classification, based on these similarity scores. We used the Python scikit-learn package to calculate ROC and AUC.

We repeated these calculations for each one of the 30 batches and calculated the mean and std of AUCs. This was done for the bird-species images and for the individual-bird images, using all DCNNs. In addition to AUCs we calculated Representation Dissimilarity Matrices (RDMs) ^52^ for images of bird-species and individual-birds using the cosine similarities between embeddings generated by the different DCNNs. For the RDMs we randomly selected 10 images out of the 50 from each class that were used for AUC calculations.

## Acknowledgments

We would like to thank Noam Avidor, Amit Bardos, Koby Boyango and Danielle Chason who performed initial training of DCNNs for bird species in earlier stages of this research as part of their undergraduate research seminar. This research was supported by a grant from the Israeli Science Foundation (ISF 917/21) to GY.

## Footnotes

1 implementation and weights from https://github.com/TreB1eN/InsightFace_Pytorch

2 Freezing a convolution layer means freezing up to the pooling operation

